## Supplemental figures for "Synaptic Potentiation of Engram cells is Necessary and Sufficient for Context Fear Memory"

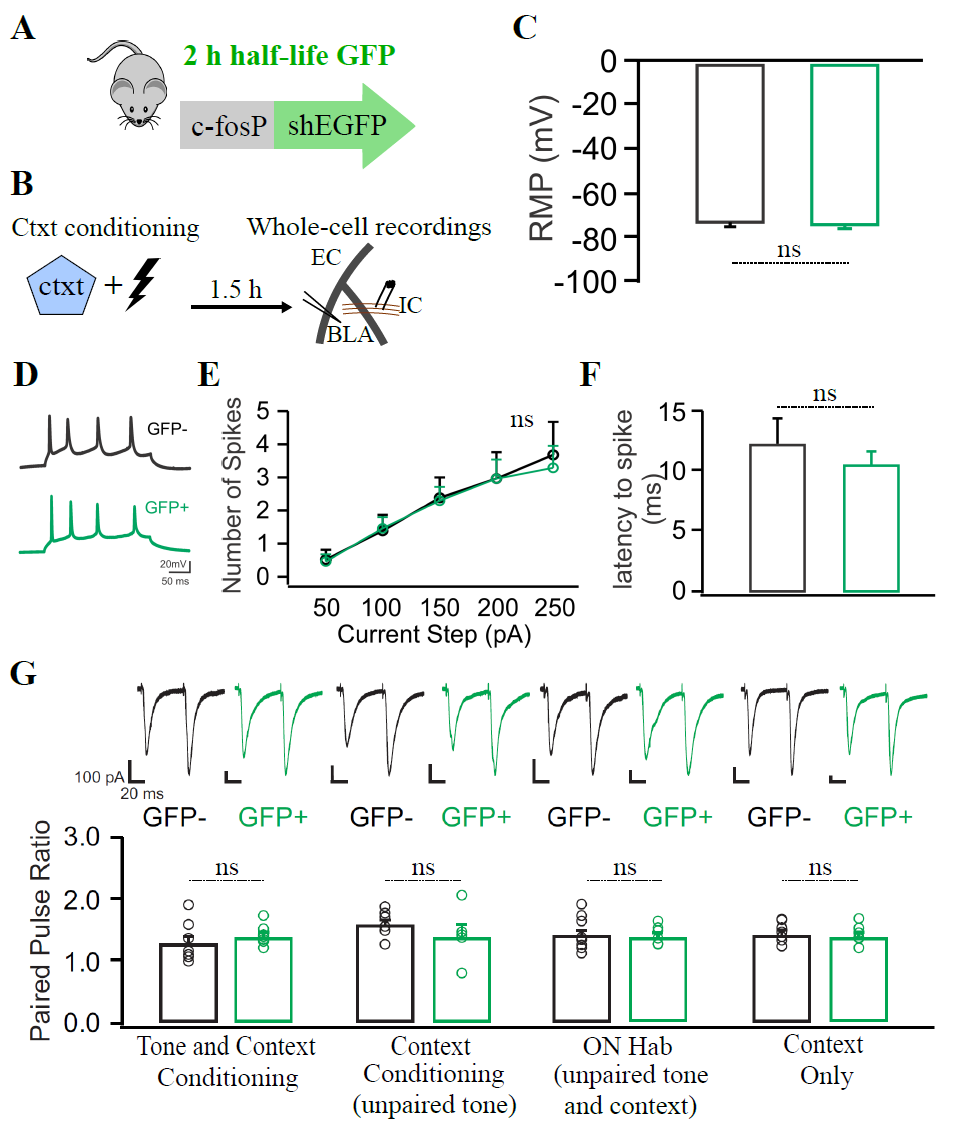

**Figure 1-figure supplement 1. cfos+ neurons tagged during training do not differ in intrinsic excitability, resting membrane potential or paired pulse ratio.** (**A**) cfos-shEGFP mouse line, which expresses a short half-life(sh) EGFP under the control of the cfos promoter. (**B**) Experimental design, in which the cfos-shEGFP mouse was used for whole-cell recordings after context conditioning. (**C**) The resting membrane potential (RMP) of GFP+ (green) and GFP- (black) neurons are similar. (**D**) Example of evoked train of action potentials with 250 pA current injection. (**E**) Both GFP+ and GFP- neurons fire similar number of action potentials in response to current injections delivered at the internal capsule. (**F**) The latency to spike between GFP+ and GFP- neurons is also similar. N = 6-7 neurons per group in (C) to (F). (**G**) The paired-pulse ratio of GFP+ and GFP- neurons show no evidence for significant presynaptic plasticity across the behavior groups tested in Fig. 1C. N =6-9 neurons per group. Unpaired t tests [(C), (E) to (G)], ns = not significant. Graph bars show mean +/- SEM.

**Figure 1-figure supplement 2. GFP expression among groups in Figure 1; Long-lasting increase in AMPAR/NMDAR ratio in Arc+ neurons tagged during training is reversed by extinction and reinstated by retraining.** (**A**) Representative coronal BLA sections of the cfos-shEGFP mouse, showing eGFP expression after Context Conditioning and Context Only exposure. Scale bar: 25 μm. (**B**) The percentage of GFP+ (GFP/DAPI) cells is similar among behavior groups in Fig. 1, with the exception of Context only exposure. The rightmost bar refers to Arc-tTA/TetO-H2BGFP double transgenic mouse, used in Fig. 1, F to H. N = 4-5 mice per group. (**C**) Arc-tTA/TetO-H2BGFP double transgenic mouse, in which the tetracycline transactivator (tTA) was knocked in the Arc gene and controls the expression of the long-lasting histone-bound GFP. (**D**) Experimental design, in which the Arc-tTA/TetO-H2BGFP mouse was used to tag Arc+ cells during training with a long-lasting histone-bound GFP for subsequent behavior or recordings 7 days later. Animals submitted to context extinction were exposed daily for 5 days to conditioned context for 30 min/day. (**E**) Freezing levels across the extinction trials and after retraining, which restores freezing to levels compared to animals that did not undergo extinction. Green: context conditioning group; black: extinction group; purple: Retraining group. N = 8 mice per group. **(F)** Extinction reverses long lasting increase in AMPAR/NMDAR ratio of learning-activated neurons (GFP+), while re-training reinstates the increase in AMPAR/NMDAR ratio in the same population of neurons activated during the original conditioning. The data for the context conditioning group is a reproduction of Fig 1H, to allow easier comparison with other groups. N = 6-9 neurons per group. ****P < 0.0001, **P < 0.01, *P < 0.05, one-way ANOVA with Tukey test (B), unpaired t test (F). Graph bars show mean +/- SEM.

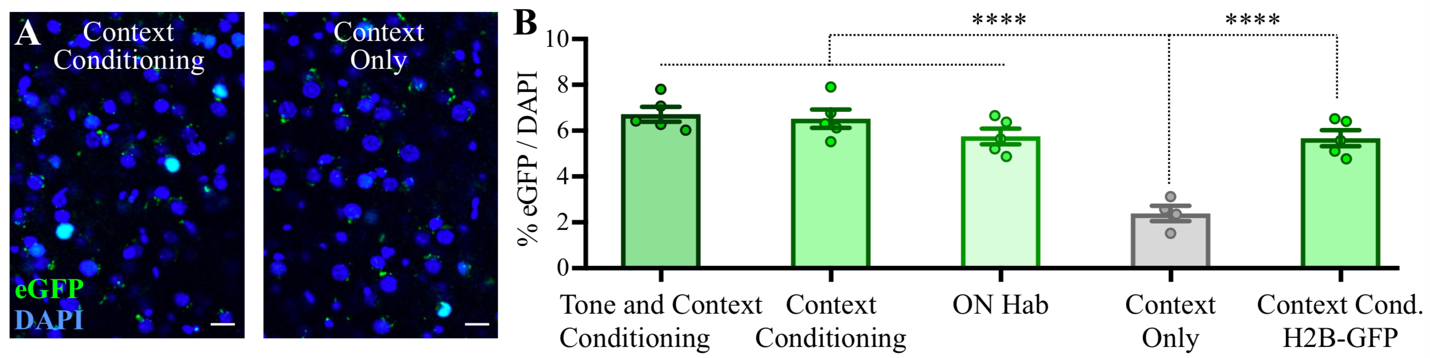

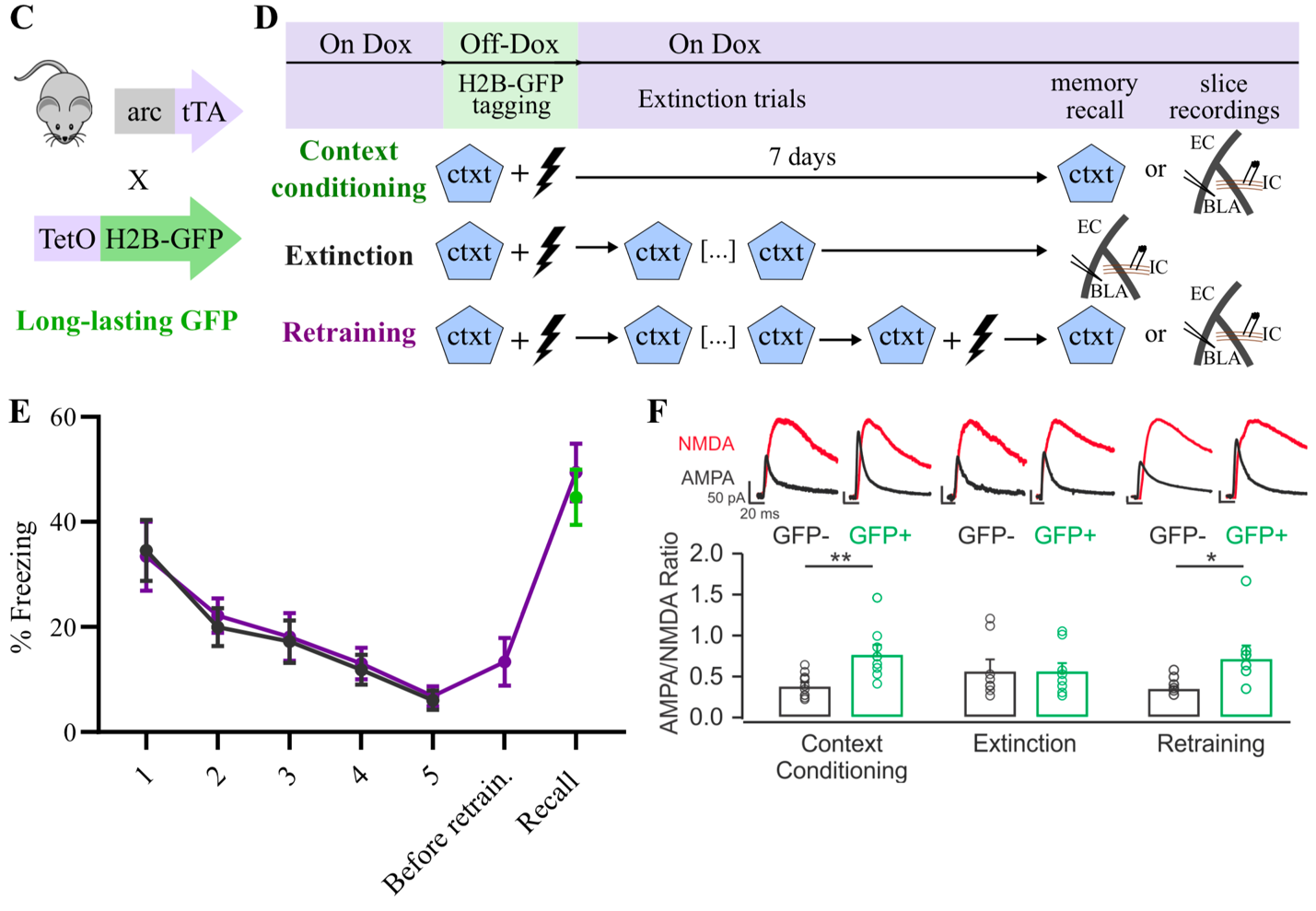

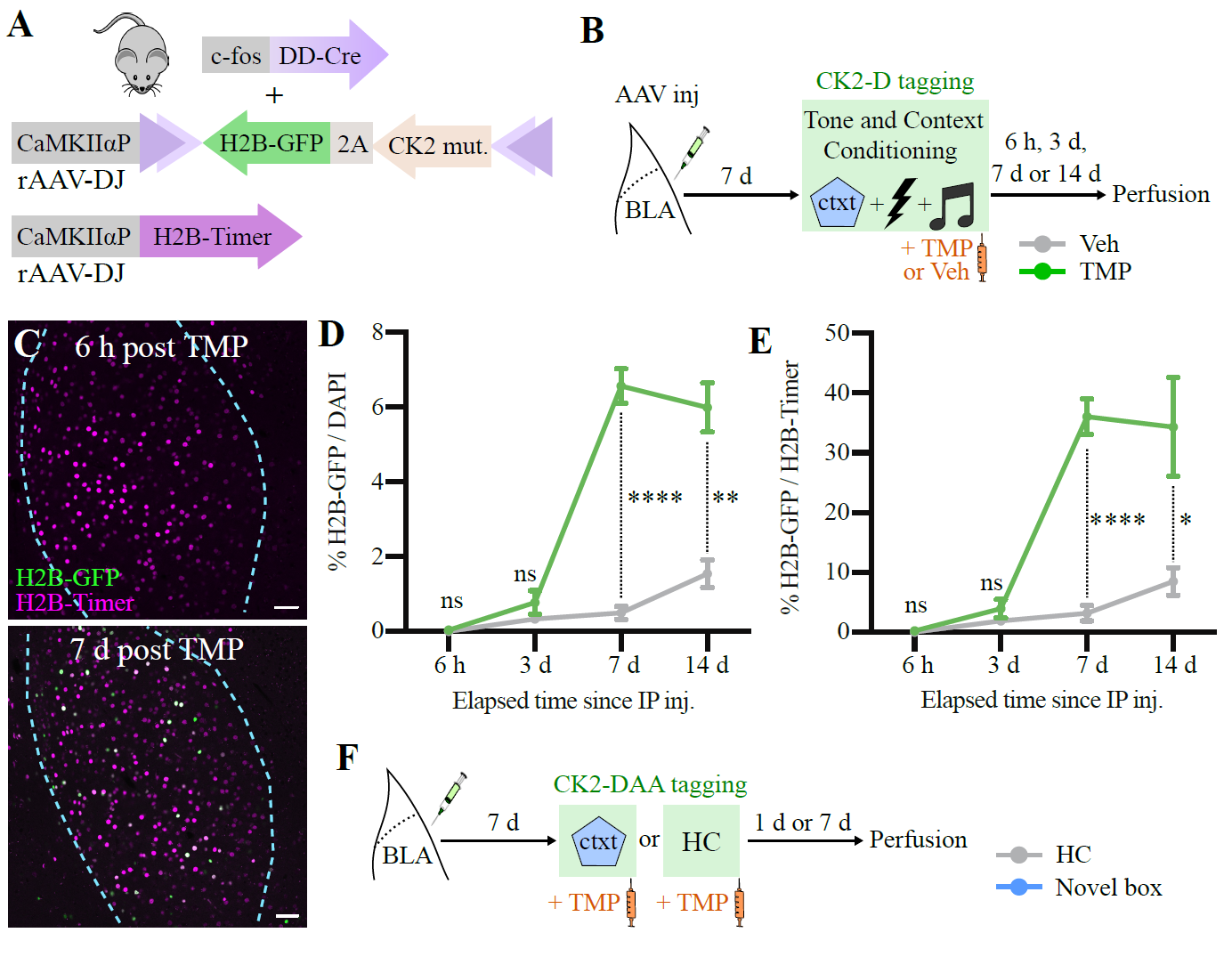

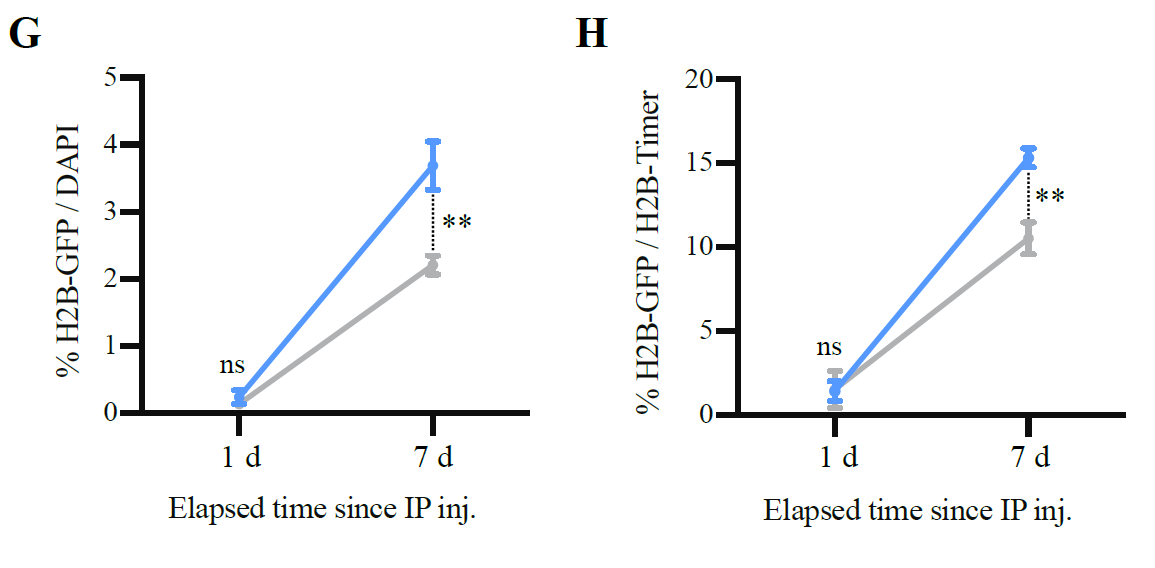

**Figure 2-figure supplement 1. Expression time course of CaMKIIα mutants** (**A**) rAAV-DJs carrying a DIO-CK2 mut. construct and a red marker (H2B-Timer) construct were co-injected bilaterally in the BLA of FDC mice. (**B**) Experimental design to determine the time course of CamKIIα-T286D (CK2-D) expression after tagging during fear conditioning. (**C**) Representative images of coronal BLA sections 6 hours and 7 days after fear conditioning followed by IP injection of TMP. BLA is highlighted within dashed cyan lines. Scale bar: 50 μm. (**D**) The percentage of total BLA cells (DAPI) that express CK2-D (H2B-GFP) after TMP or Veh injections reaches its peak 7 days after TMP injection. (**E**) A similar result is found normalizing the number of CK2-D cells (H2B-GFP) by the number of infected cells (H2B-Timer), which takes into account injection coverage variation. N = 4 mice per group per time point, except for 6 hours, where N = 3 (D) and (E). (**F**) Experimental design to determine the time course of CamKIIα-T286D/T305A/T306A (CK2-DAA) after exposure to novel box or homecage (HC). (**G**) The percentage of total BLA cells (DAPI) that express CK2-DAA (H2B-GFP) after novel box exposure is not different from homecage control 1 day after tagging. (**H**) A similar result is found normalizing the number of CK2-DAA cells (H2B-GFP) by the number of infected cells (H2B-Timer), which takes into account injection coverage variation. N = 4 mice per group per time point. ****P < 0.0001,**P < 0.01, *P < 0.05, ns, not significant, unpaired t tests. Graph bars show mean +/- SEM.

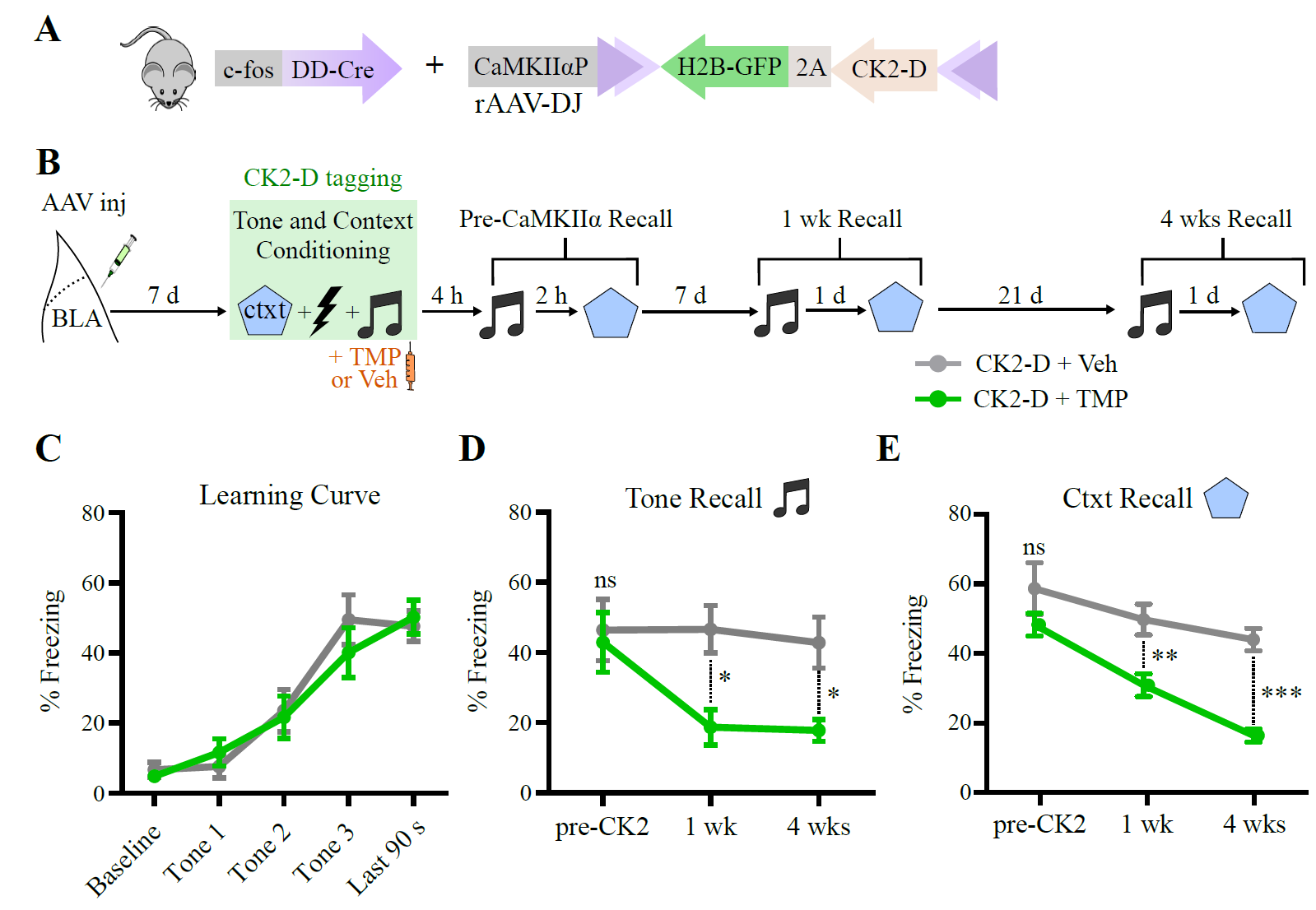

**Figure 2-figure supplement 2.** **CaMKIIα-T286D (CK2-D) effect in cfos+ BLA neurons tagged during training is long-lasting.** (**A**) rAAV-DJ carrying a Cre-dependent CK2-D-2A-H2BGFP construct was injected bilaterally in the BLA of FDC mice. (**B**) Experimental design to test the effects of CK2-D expression over time in cfos+ BLA neurons tagged during fear conditioning. (**C**) Freezing response across training trial, with three tone-shock pairings. As expected, the freezing to the tone increased after pairings with shock. (**D**) and (**E**) CK2-D mediated-memory impairment for tone (D) and context (E) lasts for at least 4 weeks. N = 7 mice per group. *P < 0.05, **P < 0.01, ***P < 0.001, ns, not significant, two-way RM ANOVA with Bonferroni test [(C) to (E)]. Graph bars show mean +/- SEM.

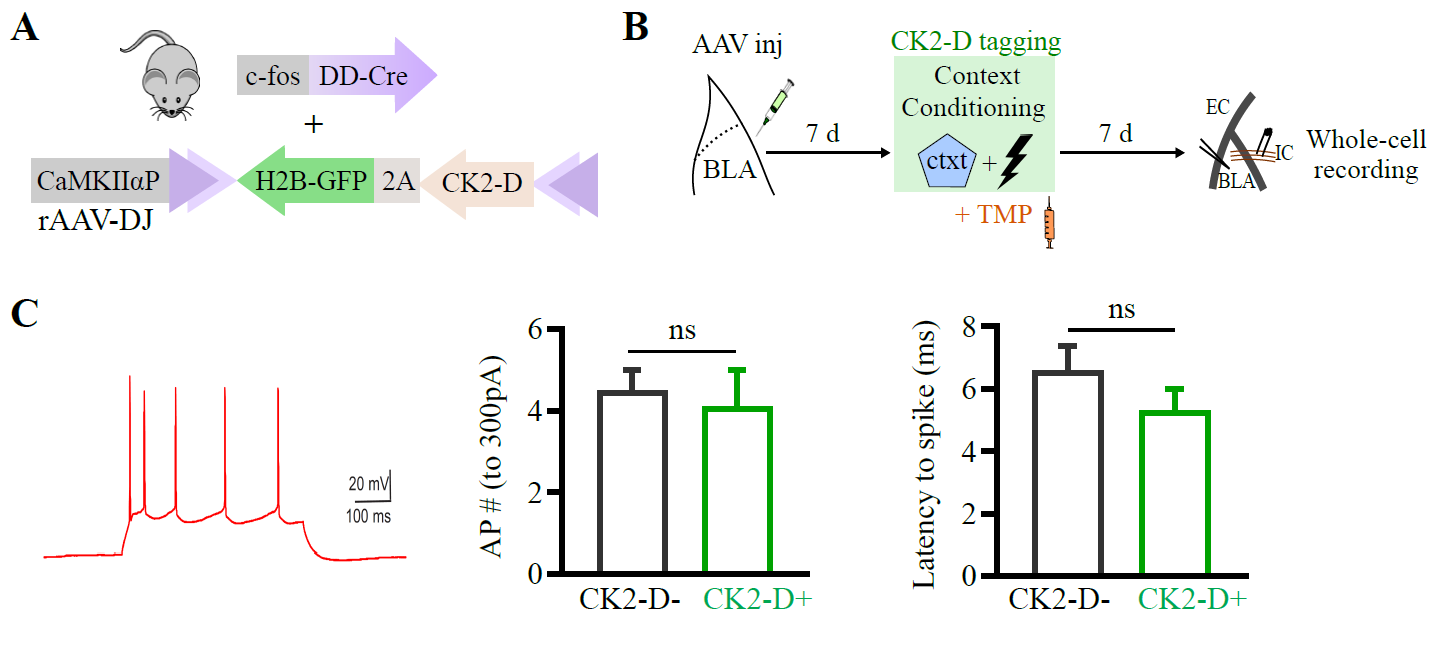

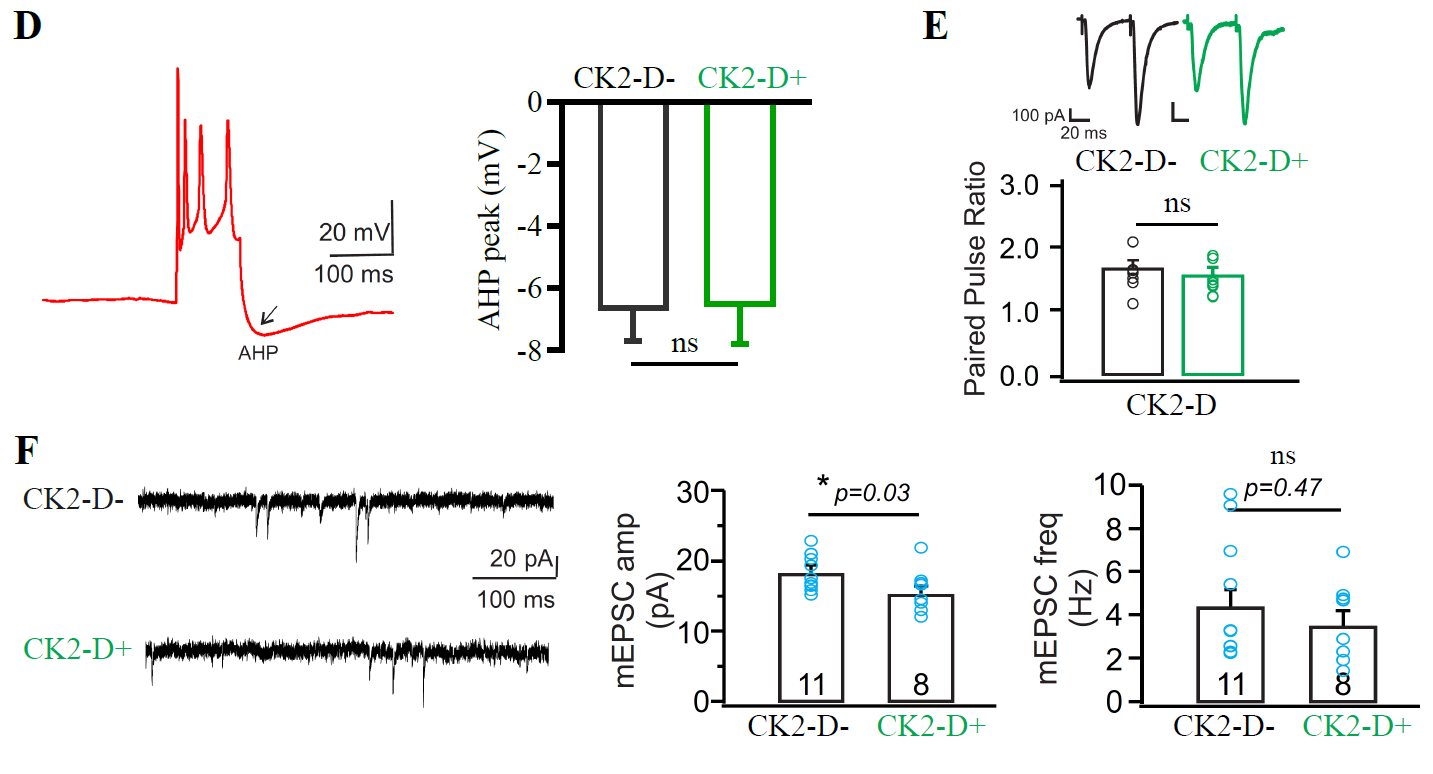

**Figure 2-figure supplement 3.** **C****aMKIIα-T286D (CK2-D) expression in cfos+ BLA neurons tagged during training affects the amplitude of mEPSCs but not the frequency, without altering the intrinsic excitability or paired-pulse ratio.** (**A**) rAAV-DJ carrying a Cre-dependent CK2-D-2A-H2BGFP construct was injected bilaterally in the BLA of FDC mice. (**B**) Experimental design to test the effects of CK2-D expression on the intrinsic excitability, paired-pulse ratio and mEPSCs of cfos+ BLA neurons tagged during context conditioning. (**C**) and (**D**) CK-2D expression does not affect the number of action potentials triggered by 300pA current injection in the internal capsule fibers neither the latency to spike (C), nor the after-hyperpolarization (AHP) peak after a train of 4 or more action potentials (D). N = 16 CK2-D- neurons and 10 Ck2-D+ neurons. Additional measurements regarding these neurons can be found on table S1. (**E**) The paired-pulse ratio of CK2-D+ and Ck2-D- neurons show no evidence for significant presynaptic plasticity after training. N = 7-8 per group. (**F**) mEPSCs from CK2-D+ neurons have smaller amplitudes, but no difference in frequency, compared to Ck2-D- neurons, suggesting post-synaptic depression. Number of recorded mEPSCs indicated within bars. *P < 0.03, ns, not significant, unpaired t tests. Graph bars show mean +/- SEM.

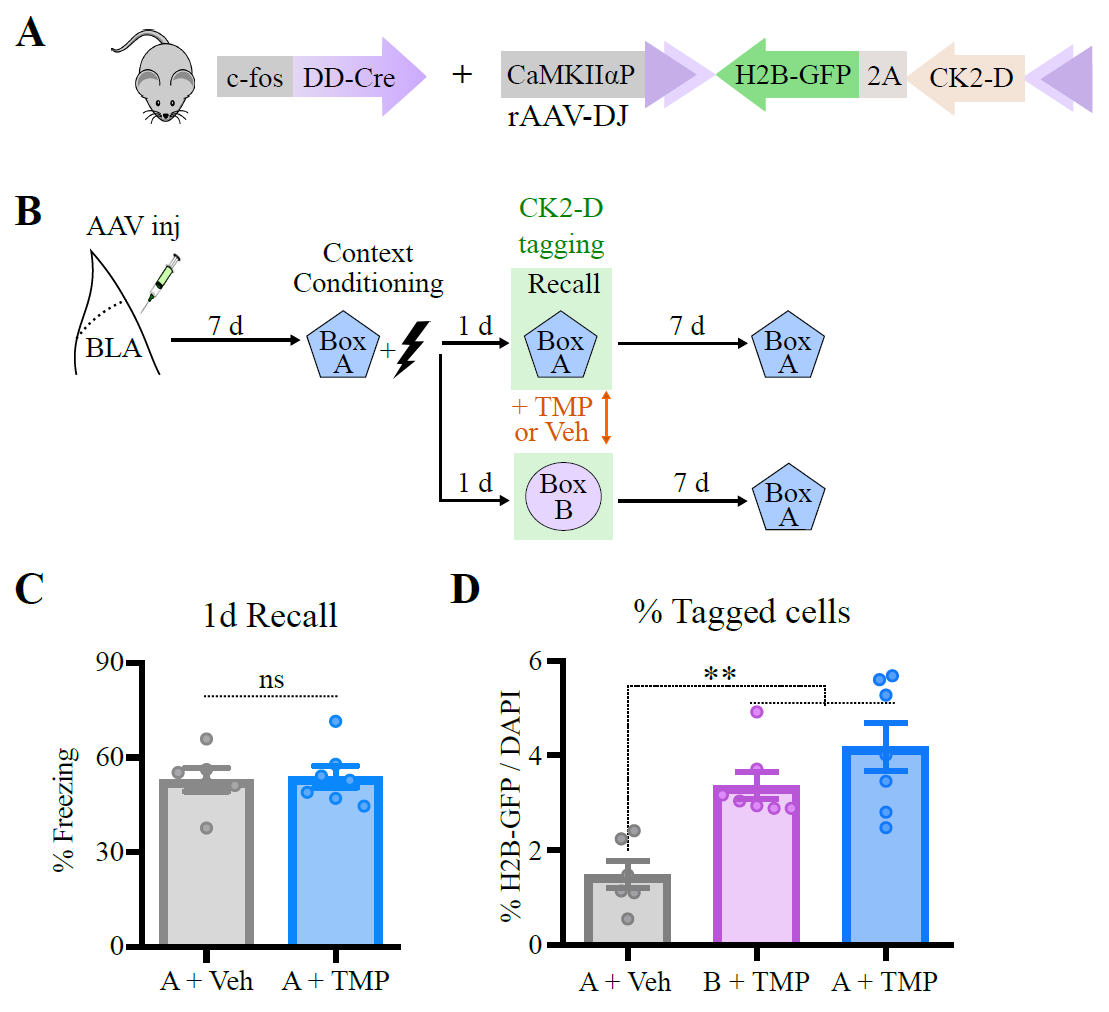

**Figure 2-figure supplement 4. Memory was intact before CaMKIIα-T286D (CK2-D) induction, and the percent of tagged cells was similar to control group, tagged during exposure to novel box (box B).** (**A**) rAAV-DJ carrying a Cre-dependent CK2-D-2A-H2BGFP construct was injected bilaterally in the BLA of FDC mice. (**B**) Experimental design to test specificity of effect to the tagged context. (**C**) 1-day (1d) recall in box A before CK2-D induction shows that memory was similar between groups (compare with Figure 2K, after CK2-D induction). (**D**) Percent of tagged cells (H2B-GFP+) was similar between group tagged during recall to conditioning box (box A) and exposure to novel box (box B). N = 6 mice per group. **P < 0.01, ns, not significant, unpaired t test (C), or one-way ANOVA with Tukey test (D). Graph bars show mean +/- SEM.

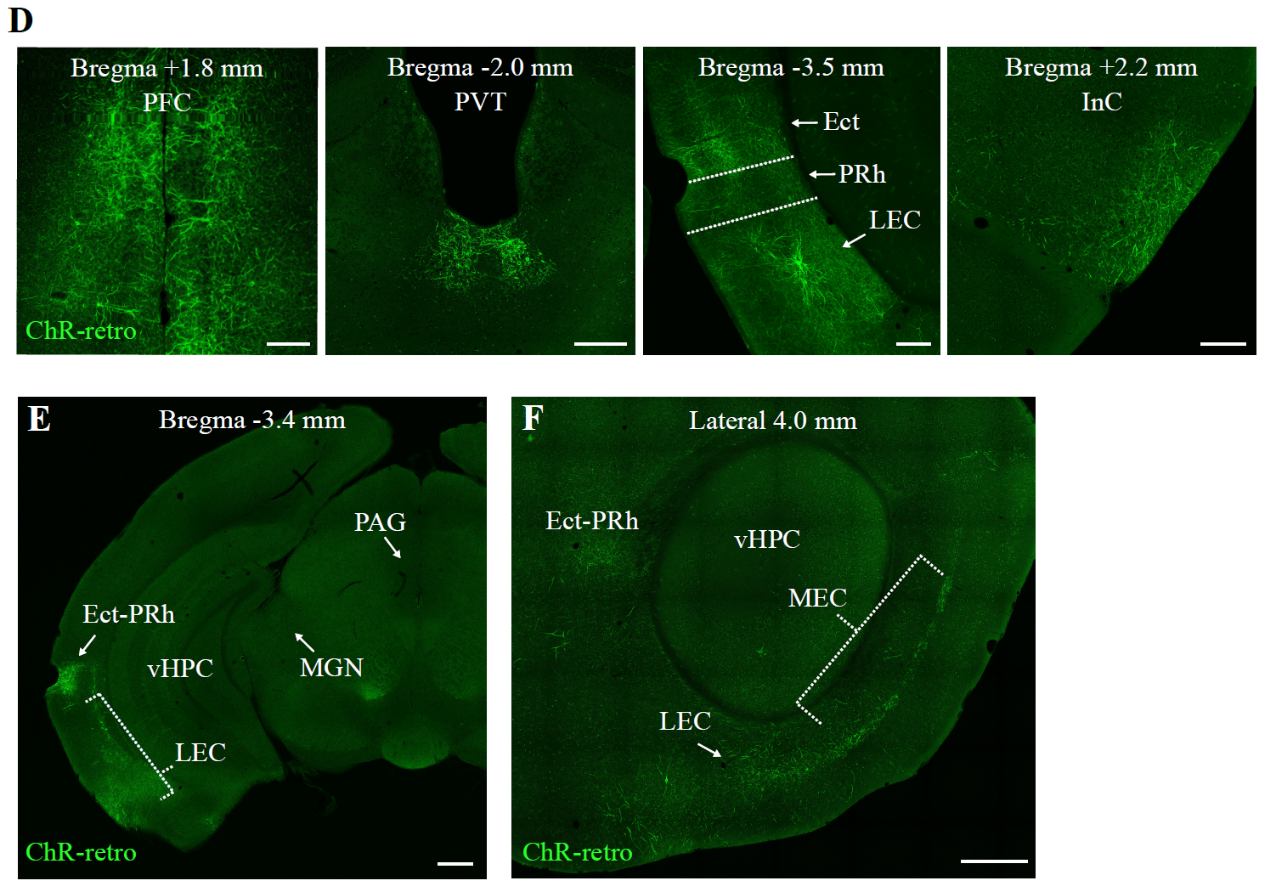

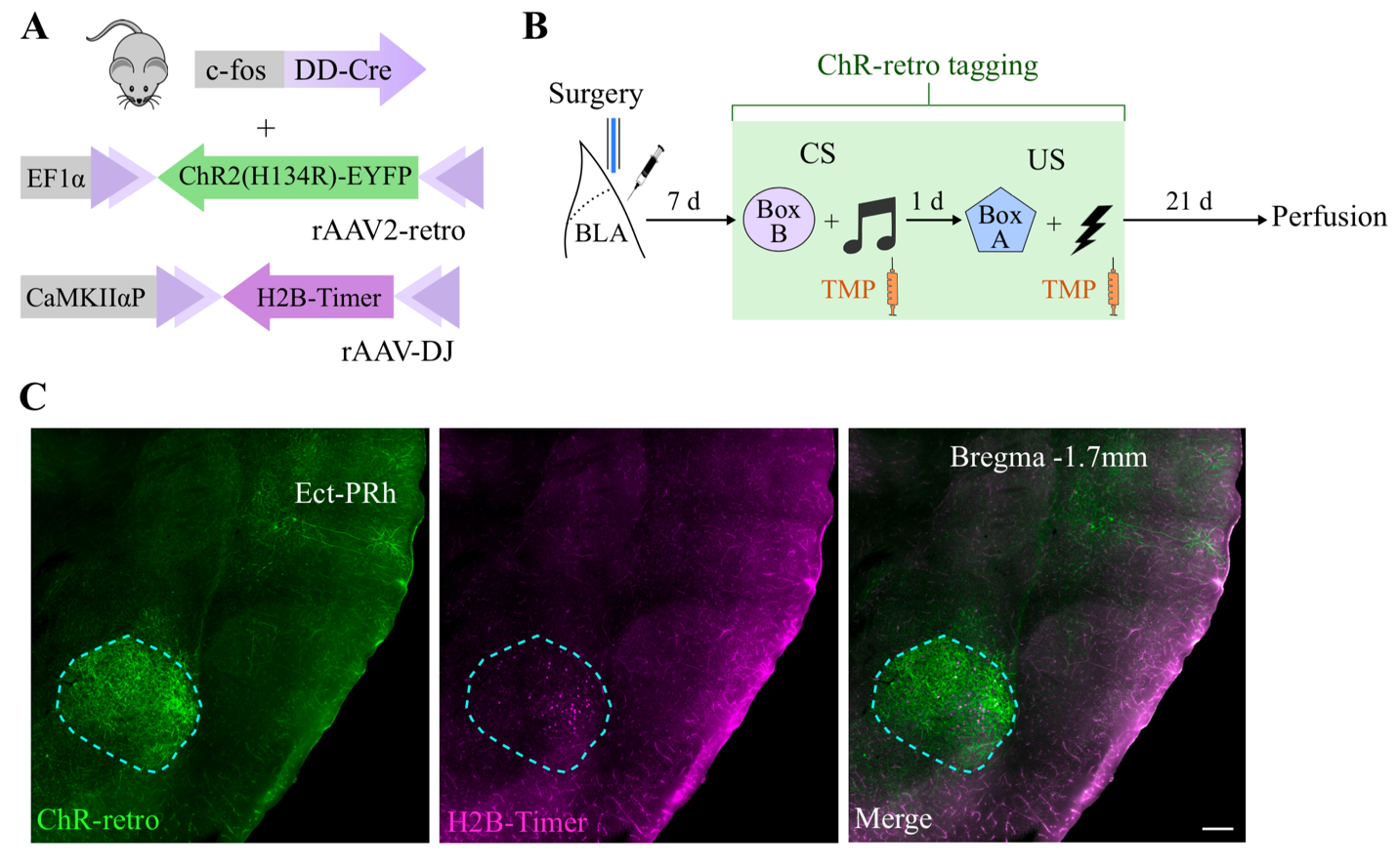

**Figure 3-figure supplement 1****. ChR-retro expression pattern after injection in BLA.** (**A**) rAAV2-retro carrying a DIO-ChR2(H134R)-eYFP (ChR-retro) construct was billaterally co-injected with a non-retrograde rAAV-DJ carrying a H2B-Timer construct in the BLA of FDC mice (**B**) Experimental design to investigate the ChR-retro spread after injection in the BLA. (**C**) Coronal BLA section showing that H2B-Timer expression is restricted to the injection target, while ChR-retro spreads to BLA inputs, such as Ect-PRh. Scale bar: 200 μm. (**D**) Coronal sections showing the main areas where ChR-retro expression was detected. Scale bar: 200 μm. (**E**) Coronal section highlighting regions that do not show substantial ChR-retro expression, such as vHPC, MGN and PAG. Scale bar: 500 μm. (**F**) Sagittal section highlighting ChR-retro expression at MEC, as well as LEC and Ect-PRh. Scale bar: 500 μm. BLA: Basolateral amygdala; Ect-PRh: Ectorhinal and Perirhinal cortices; InC: Insular cortex; LEC: Lateral Entorhinal cortex; MEC: Medial Entorhinal cortex; MGN: medial geniculate nucleus; PAG: periaqueductal gray; PFC: Prefrontal cortex, including prelimbic, infralimbic and mediorbital regions; PVT: Paraventricular nucleus of thalamus, vHPC: ventral hippocampus.

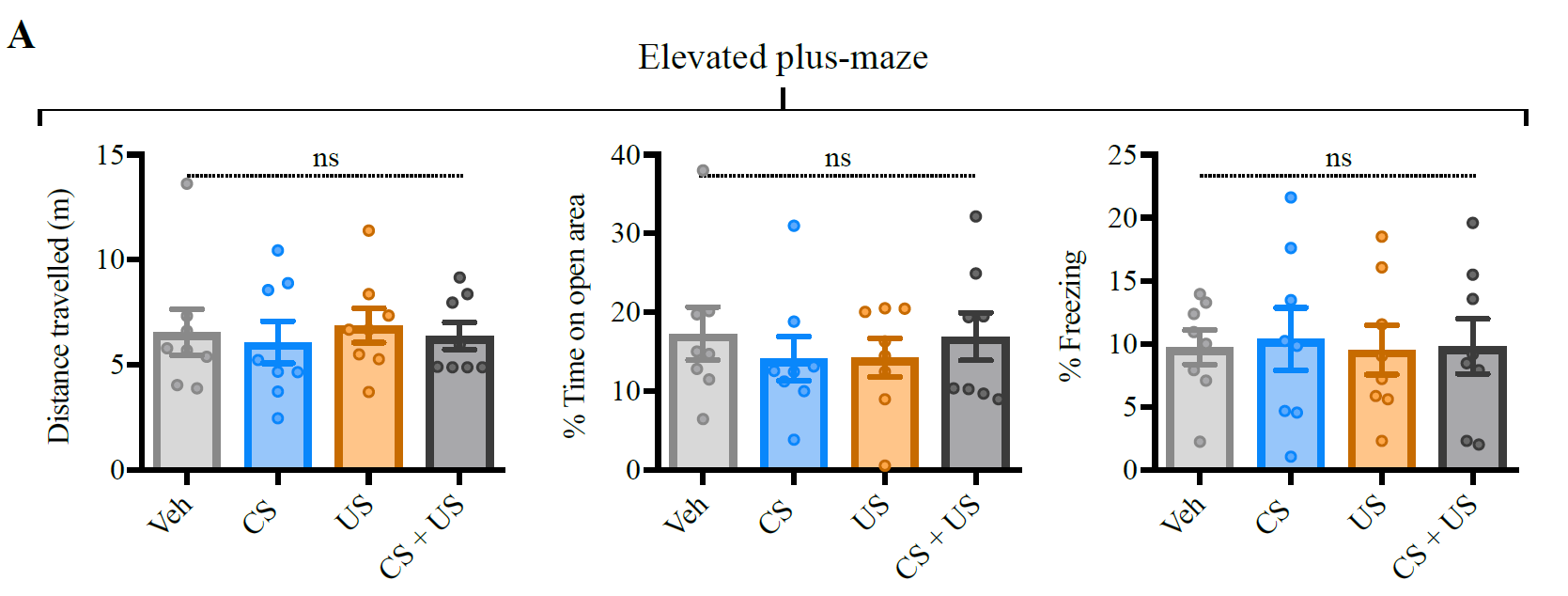

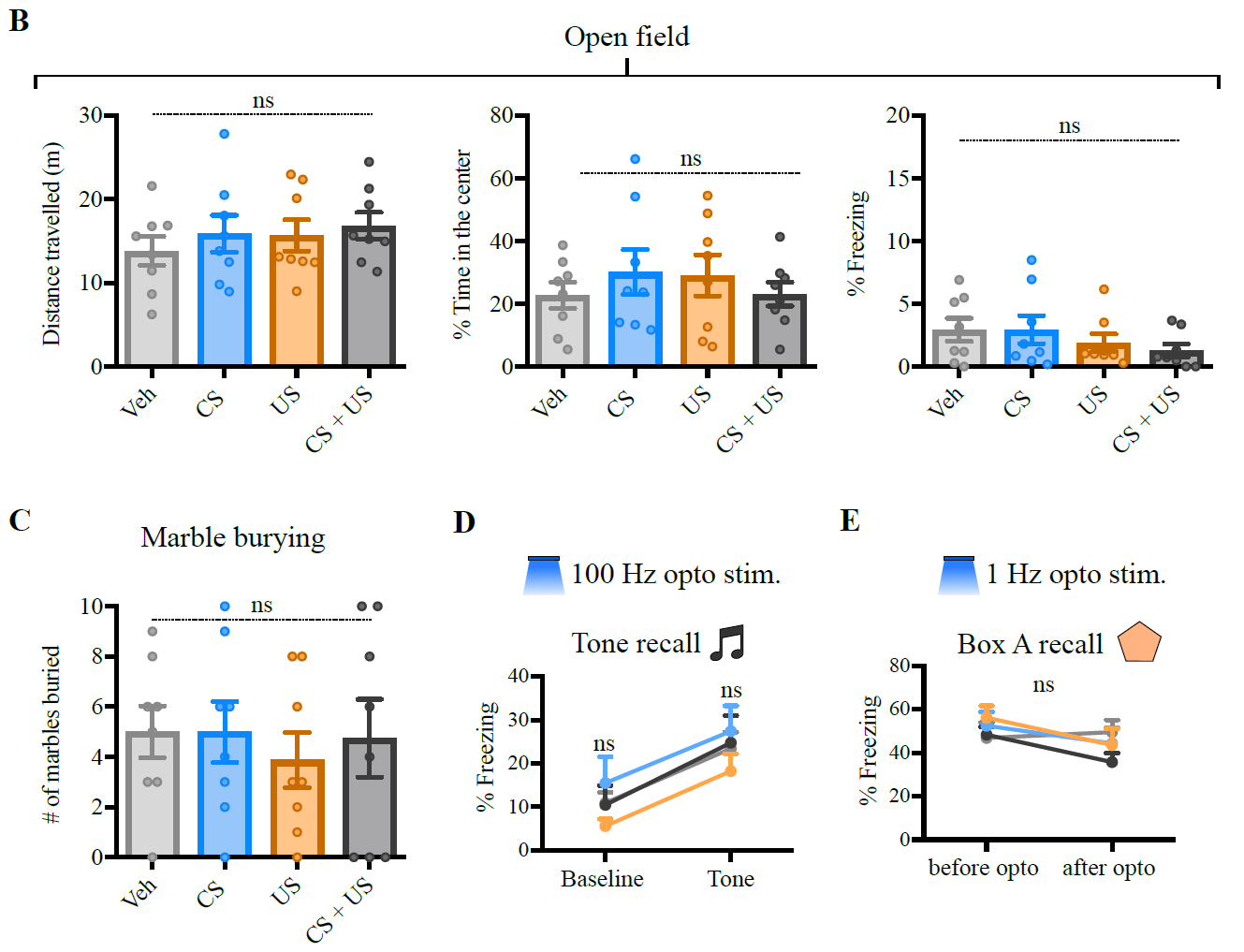

**Figure 3-figure supplement 2. 100 Hz optogenetic stimulation of CS and/or US inputs in the BLA does not interfere with mouse behavior in anxiety tests, nor induces tone conditioning, while 1 Hz optogenetic stimulation does not affect memory to box A.** (**A**) and (**B**) No change observed between control (Veh) and other groups on Elevated plus-maze (A) or Open field (B) at distance travelled (left), time on open area/center of the field (center) or freezing (right). (**C**) The number of marbles buried in the Marble burying test were also similar across groups. (**D**) No freezing to tagged tone was observed after 100 Hz optogenetic stimulation perhaps due to insufficient labeling of auditory inputs. (**E**) 1 Hz optogenetic stimulation did not affect natural memory to box A. N = 8 mice per group. These tests are part of a large experiment, which is summarized in Fig. 3C. Ns, not significant. One-way ANOVA [(A) to (C)] or two-way RM ANOVA [(D) and (E)]. Graph bars show mean +/- SEM.

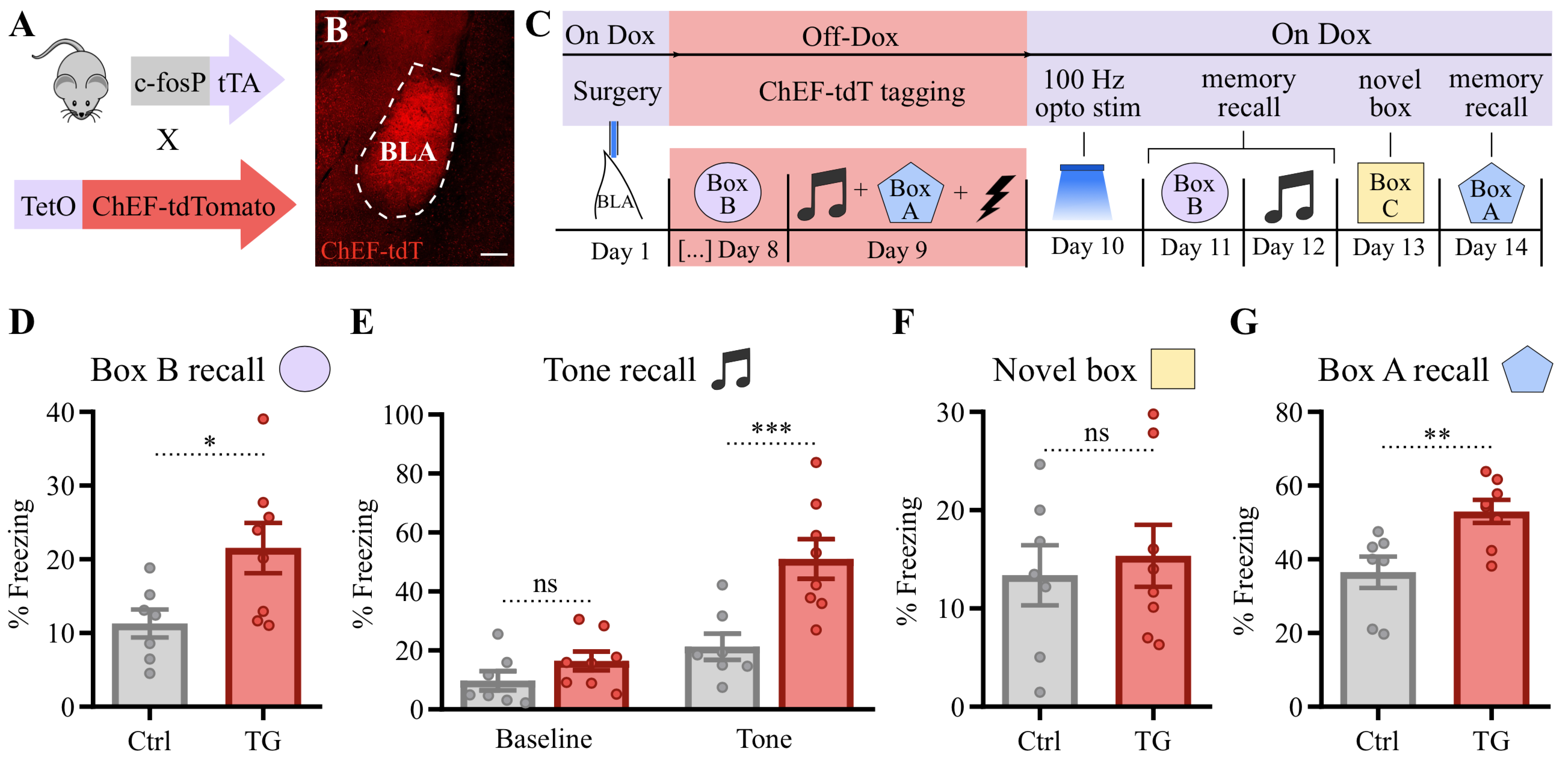

**Figure 3-figure supplement 3. 100 Hz optogenetic stimulation of CS and US inputs in the BLA creates *de novo* aversive associative memories in transgenic mouse expressing ChEF (a ChR variant) in tagged cfos+ neurons.** (**A**) Double transgenic mouse expressing the Channelrhodopsin variant ChEF fused to tdTomato (tdT) under the control of the Tet system driven by the cfos promoter. (**B**) Coronal section at Bregma -1.2 mm showing ChEF-tdT expression in the BLA one day after tagging shown in (C). Scale bar: 200 μm. (**C**) Experimental design used to test whether 100 Hz optogenetic stimulation of CS and US inputs in the BLA may create a de novo memory association. (**D**) to (**G**) 100 Hz light stimulation induced a de novo fear memory to box B (D) and to the tone (E) (unpaired with the shock during tagging). Effect was context specific, as no enhanced freezing to a novel box was detected (F). The natural fear memory to box A was potentiated by 100 Hz light stimulation (G). Control group (Ctrl) was composed of single transgenic mice from the same cohort of the double transgenic group (TG). N = 7-8 mice per group. ***P < 0.0001 **P < 0.01, *P < 0.05, ns, not significant, unpaired t test [(D), (F) and (G)], or two-way RM ANOVA with Bonferroni test (E). Graph bars show mean +/- SEM.

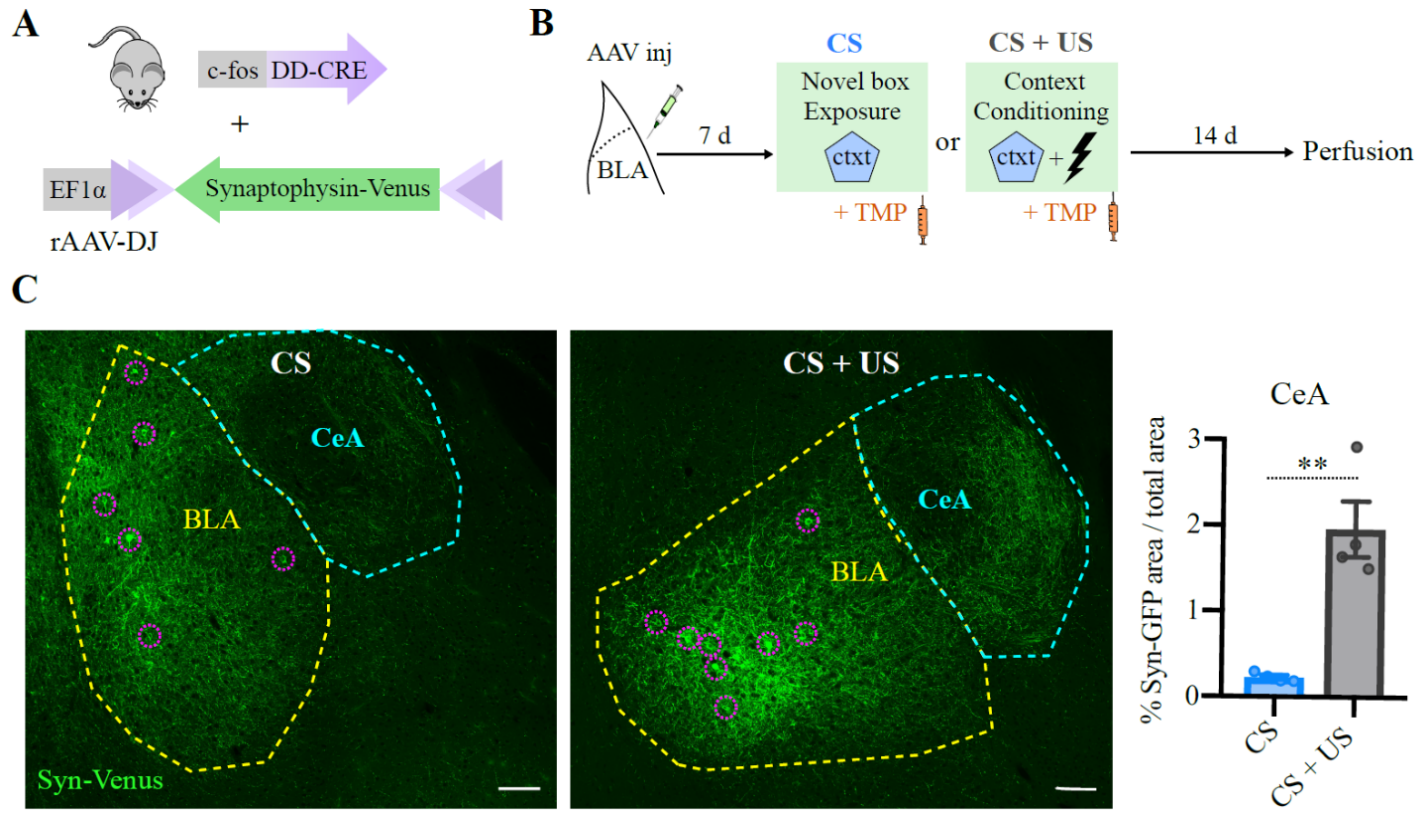

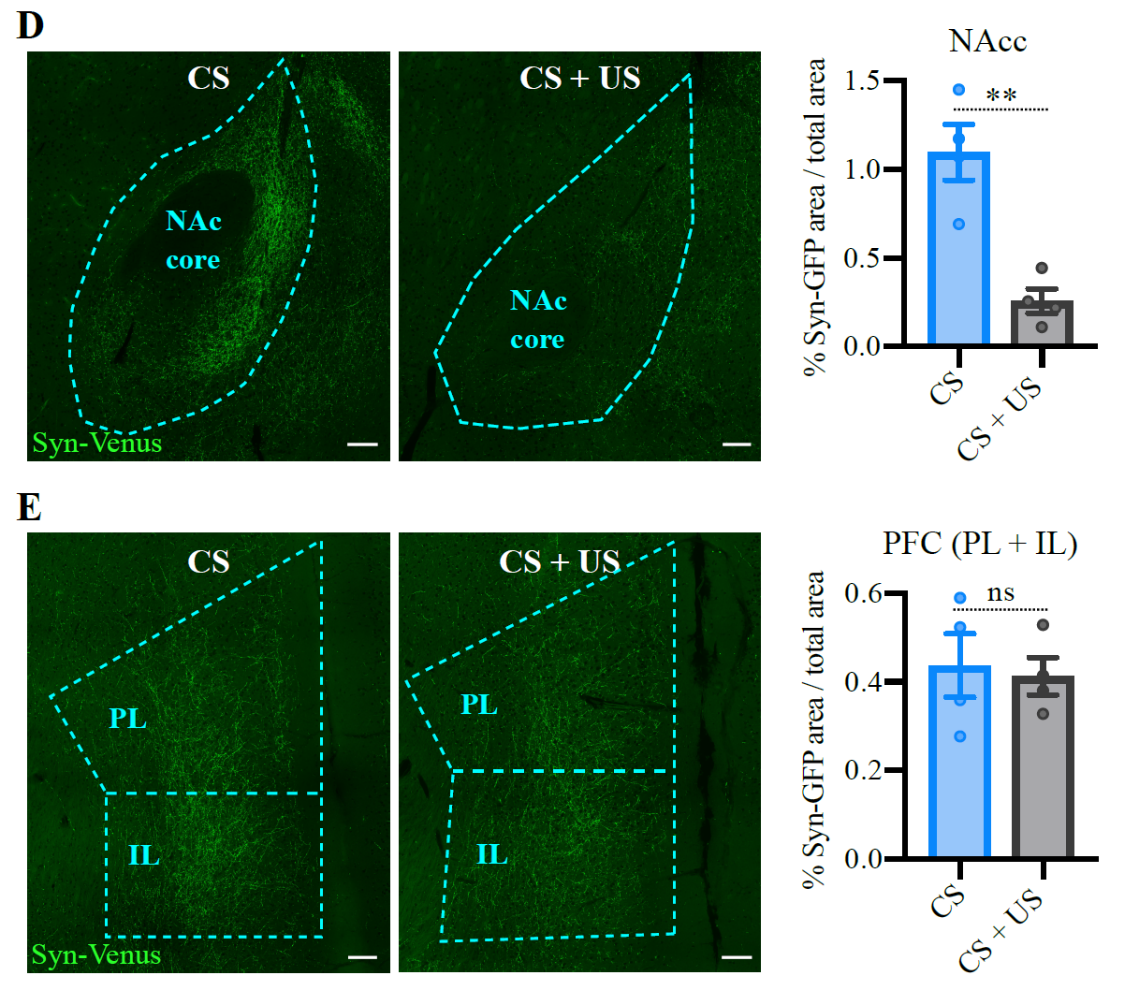

**Figure 3-figure supplement 4 . The projection targets of BLA neurons activated by context conditioning (CS+US) are biased towards the Central Amygdala.** (**A**) rAAV-DJ encoding a DIO-Synaptophysin-Venus (Syn-Venus) was unilaterally injected in the BLA of FDC mice. (**B**) Experimental design to compare the projection targets between BLA neurons activated by CS (novel box) and CS + US (context conditioning). (**C**) Representative coronal sections showing the injection site at BLA (yellow dashed line), with visible targeted neurons highlighted in purple dashed circles, and the projections to the central amygdala (CeA), which are stronger in the CS + US group (center), quantified in the graph on the right. (**D**) Representative coronal Nucleus Accumbens core (NAcc) sections showed the opposite trend, with stronger projections in the CS group, quantified in the graph on the right. (**E**) Representative coronal pre-frontal cortex sections, including prelimibic (PL) and infralimbic (IL) regions, where no substantial difference between the groups was found, quantified in the graph on the right. Scale bar: 100 μm. N = 4 mice per group. **P < 0.01, ns, not significant, unpaired t test. Graph bars show mean +/- SEM.

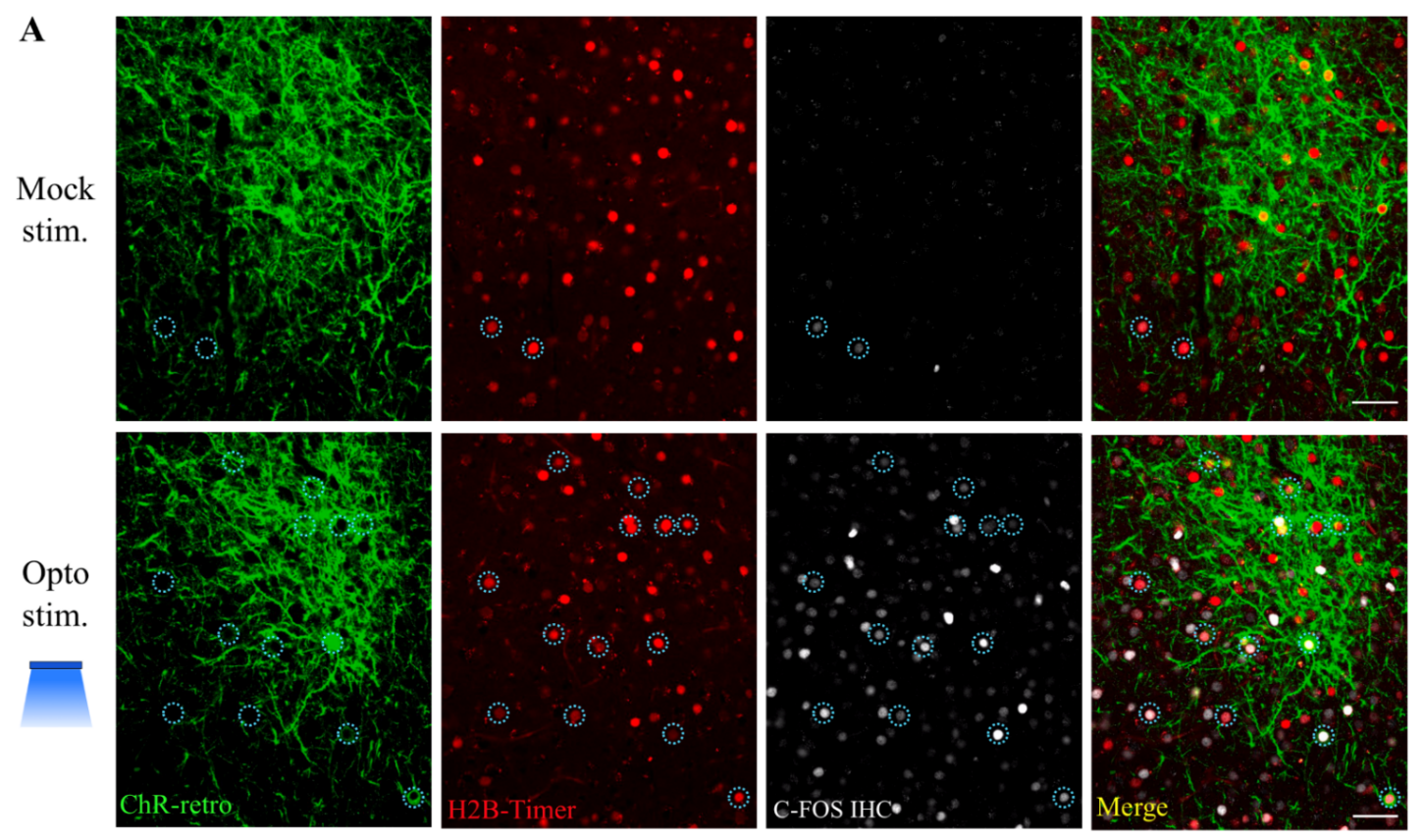

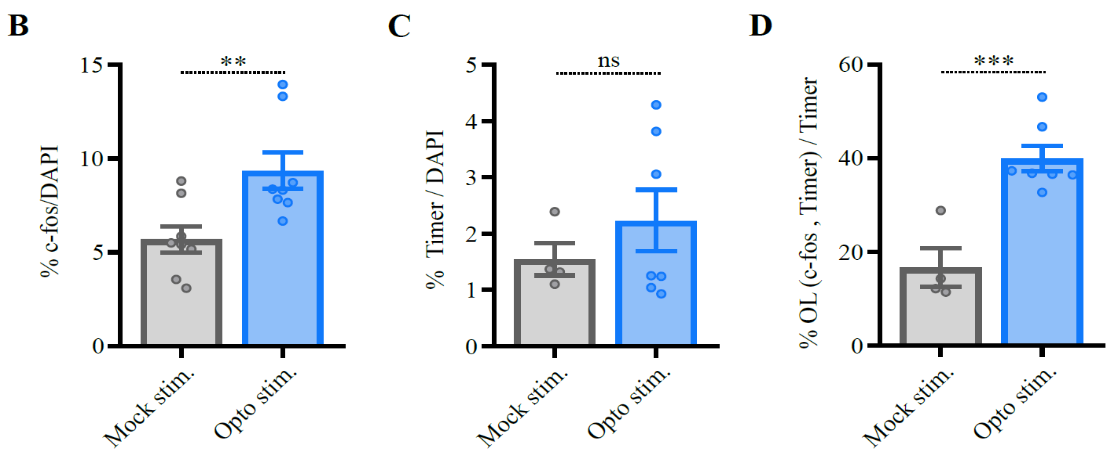

**Figure 3-figure supplement 5. 20 Hz optogenetic stimulation induces increased cfos levels in the BLA, which overlaps with tagged US neurons.** (**A**) Representative coronal BLA section showing the expression of ChR-retro and the co-injected nuclear marker H2B-Timer along with cfos immunohistochemistry (IHC) in animals perfused 90min after 20Hz opto stim. or mock stim. Dotted circles highlight tagged cells that were reactivated (both Timer+ and cfos+, quantified in D). The complete experimental design is illustrated in Figure 3C. Only animals that were tagged during US exposure, therefore exhibited freezing to 20 Hz optogenetic stimulation (Fig. 3G) were included in this experiment (“US” and “CS + US” mice were equally divided among the two groups and pooled together, as there was no significant difference between them). Scale bar: 50 μm. (**B**) 20 Hz optogenetic stimulation induced increase in overall cfos levels in BLA. N = 8 mice per group. (**C**) Timer levels were similar between groups. (**D**) Neurons that were tagged during US exposure (Timer+) tend to be reactivated (cfos+) during 20 Hz opto stim. OL: overlap. N = 4 in “mock stim.” group and 8 in “opto stim.” group in (C) and (D). ***P < 0.0001 **P < 0.01, ns, not significant, unpaired t test. Graph bars show mean +/- SEM.

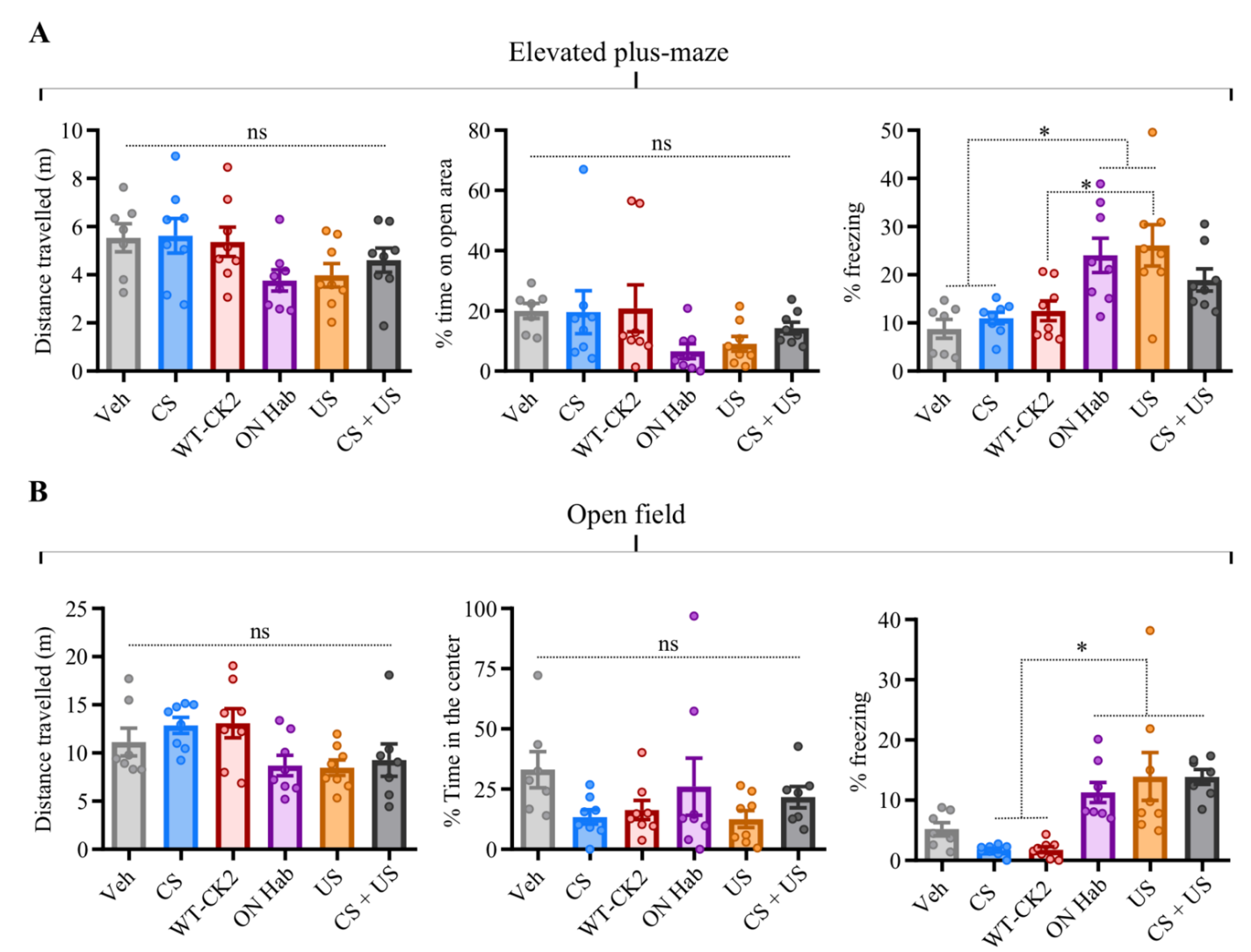

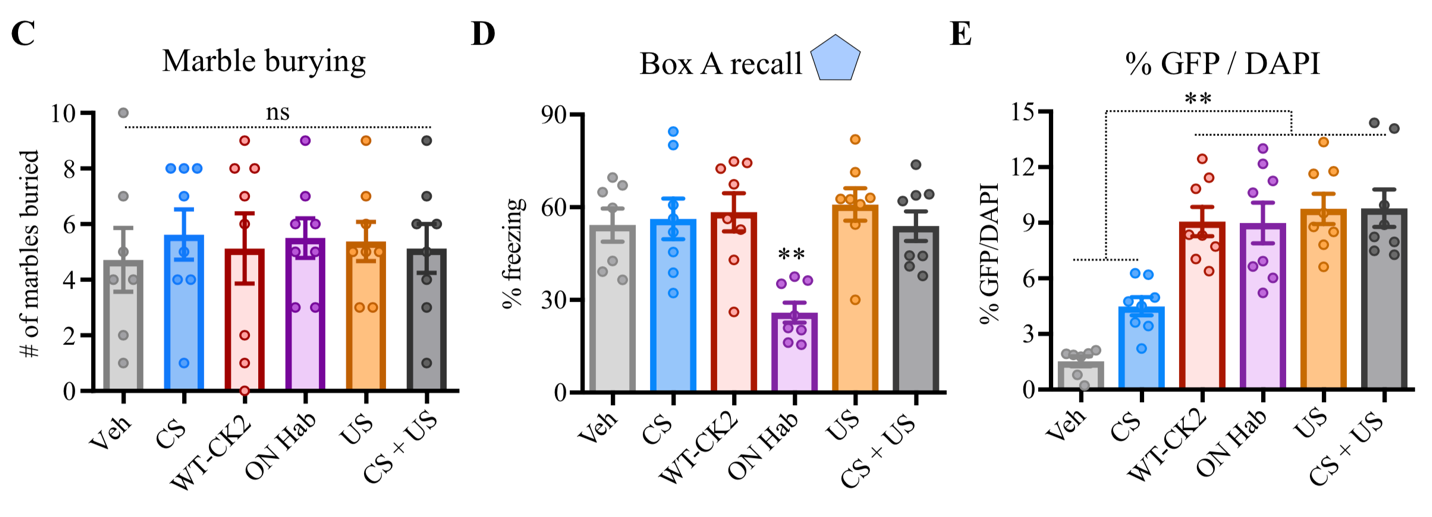

**4-figure supplement 1. CK2-DAA expression in the BLA does not interfere with locomotion or behavior in anxiety tests, despite affecting freezing.** (**A**) and (**B**) No change observed between control (Veh) and other groups on Elevated plus-maze (A) or Open field (B) at distance travelled (left) or time on open area/center of the field (center), despite some difference in freezing levels (right). (**C**) The number of marbles buried in the Marble burying test were also similar across groups. (**D)** The ON Hab group showed reduced freezing levels to box A. (**E**) The percentage of tagged cells (H2B-GFP+) were smaller in Veh and CS groups. N = 8 mice per group, except on open field, where 1 outlier animal was excluded from “CS + US group” due to elevated freezing at the center of the arena. The tests are part of a large experiment, which is summarized in Fig. 4D. **P < 0.01, *P < 0.05, ns, not significant. One-way ANOVA with Tukey test. Graph bars show mean +/- SEM.

**Table 1. CaMKIIα-T286D (CK2-D) expression in cfos+ BLA neurons tagged during training does not affect intrinsic electrophysiological properties.** Results of unpaired t tests and the respective P value is included for every corresponding measure. Part of the data described here is plotted on fig. S4. AP: action potential, V resting: resting potential, Rm: Membrane resistance, AHP: afterhyperpolarization.

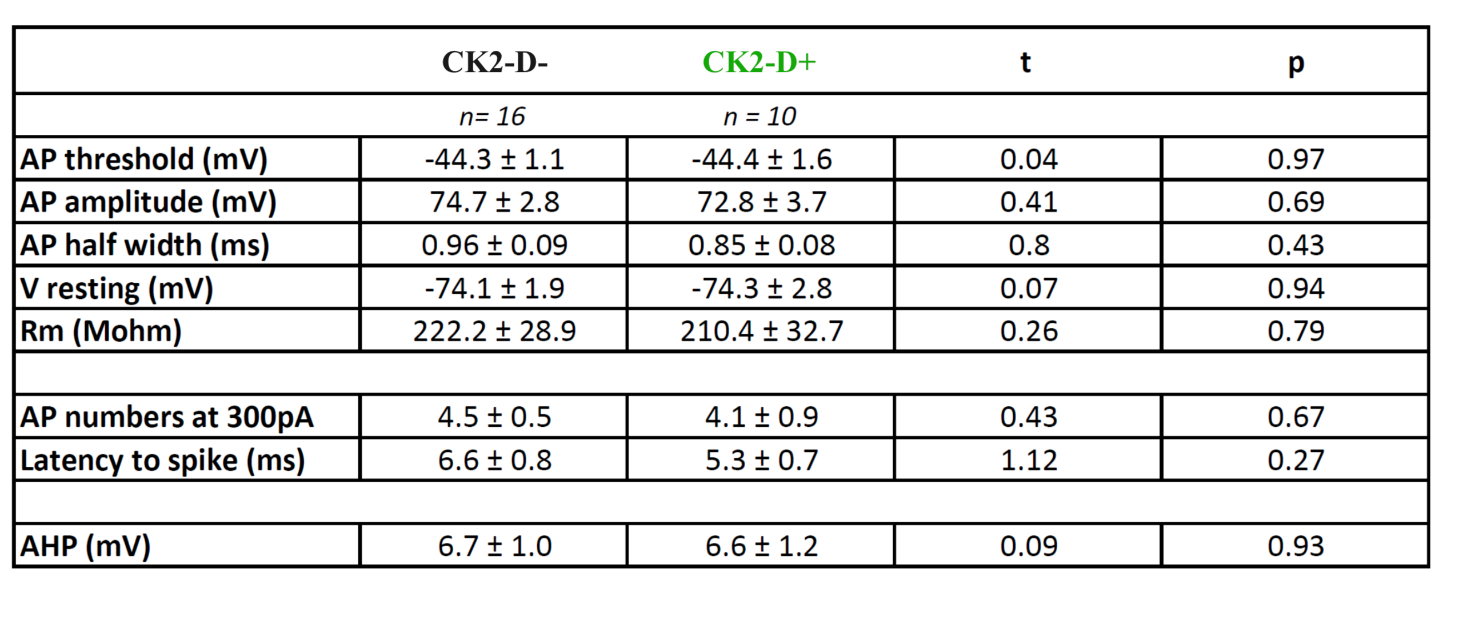

**Table 2. CaMKIIα-T286D/T305A/T306A (CK2-DAA) expression in cfos+ BLA neurons tagged during novel box exposure does not affect their intrinsic excitability.** Results of unpaired t tests and the respective P value is included for every corresponding measure. AP: action potential, V resting: resting potential, Rm: Membrane resistance, AHP: afterhyperpolarization.

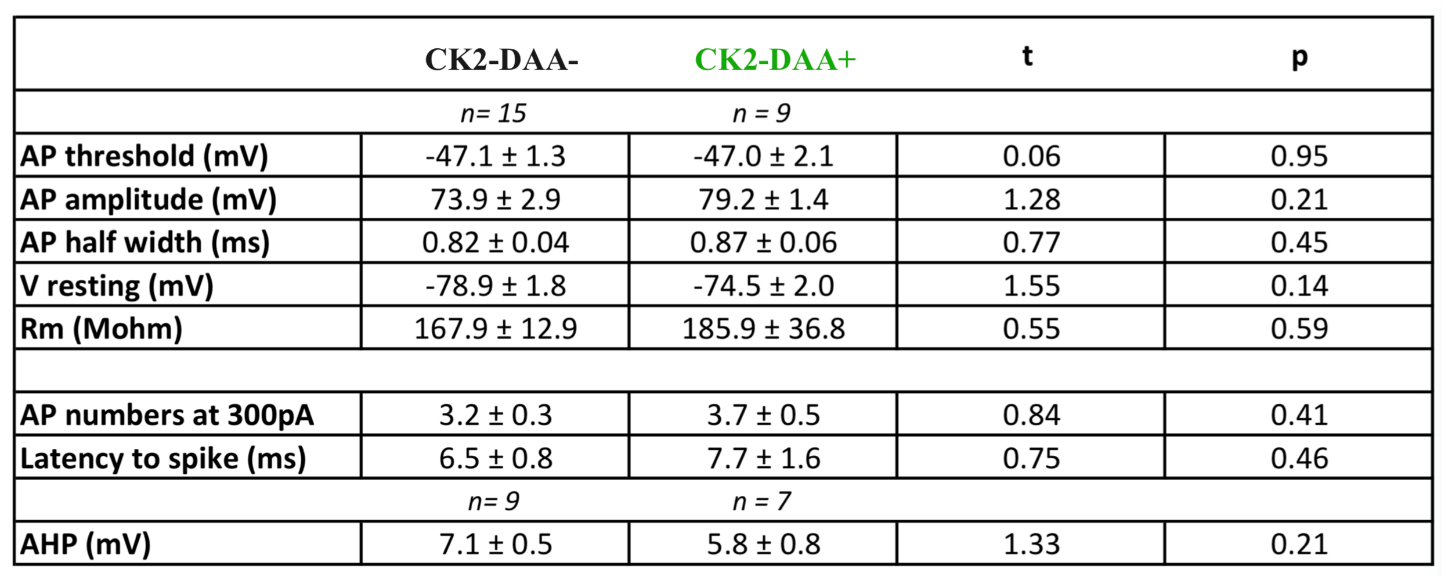
